# Distinct immune evasion in APOBEC-enriched, HPV-negative HNSCC

**DOI:** 10.1101/586396

**Authors:** Clemens Messerschmidt, Benedikt Obermayer, Konrad Klinghammer, Sebastian Ochsenreither, Denise Treue, Albrecht Stenzinger, Hanno Glimm, Stefan Fröhling, Thomas Kindler, Christian H. Brandts, Klaus Schulze-Osthoff, Wilko Weichert, Ingeborg Tinhofer, Frederick Klauschen, Ulrich Keilholz, Dieter Beule, Damian T. Rieke

**Affiliations:** Core Unit Bioinformatics, Berlin Institute of Health (BIH), 10178 Berlin, Germany; Charité – Universitätsmedizin Berlin, corporate member of Freie Universität Berlin, Humboldt-Universität zu Berlin, and Berlin Institute of Health, Berlin, Germany; Department of Hematology and Oncology, Charité – Universitätsmedizin Berlin, corporate member of Freie Universität Berlin, Humboldt-Universität zu Berlin, and Berlin Institute of Health, Berlin, Germany; Charité Comprehensive Cancer Center, Charité – Universitätsmedizin Berlin, corporate member of Freie Universität Berlin, Humboldt-Universität zu Berlin, and Berlin Institute of Health, Berlin, Germany; German Cancer Consortium (DKTK) and German Cancer Research Center (DKFZ), Heidelberg, Germany; Institute of Pathology, Charité – Universitätsmedizin Berlin, corporate member of Freie Universität Berlin, Humboldt-Universität zu Berlin, and Berlin Institute of Health, Berlin, Germany; Institute of Pathology, Heidelberg University Hospital, Heidelberg 69120, Germany; Department of Translational Medical Oncology, National Center for Tumor Diseases (NCT) Dresden and German Cancer Research Center (DKFZ), Dresden, Germany; University Hospital Carl Gustav Carus, Technische Universität Dresden, Dresden, Germany; Department of Translational Medical Oncology, National Center for Tumor Diseases (NCT) Heidelberg and German Cancer Research Center (DKFZ), Heidelberg, Germany; Department of Hematology, Medical Oncology & Pneumology, University Medical Center, Mainz, Germany; University Cancer Center Mainz (UCT), Johannes Gutenberg-University, Mainz, Germany; University Cancer Center Frankfurt (UCT), Goethe University, Frankfurt, Germany; Department of Medicine, Hematology/Oncology, Goethe University, Frankfurt, Germany; Interfaculty Institute of Biochemistry, Tübingen University, Germany; Institute of Pathology, Technical University Munich, Munich, Germany; Department of Radiooncology and Radiotherapy, Charité – Universitätsmedizin Berlin, corporate member of Freie Universität Berlin, Humboldt-Universität zu Berlin, and Berlin Institute of Health, Berlin, Germany; Max Delbrück Center for Molecular Medicine in the Helmholtz Association, Berlin, Germany; Berlin Institute of Health (BIH), 10178 Berlin, Germany

**Author notes:** **Corresponding Authors**: Dieter Beule, Core Unit Bioinformatics, Berlin Institute of Health, Berlin, Germany And Damian T. Rieke, Charité Comprehensive Cancer Center, Charitéplatz 1, 10117 Berlin.

**Keywords:** Immune checkpoint inhibition, head and neck cancer, APOBEC, mutational burden, mutational signature

## Abstract

Immune checkpoint inhibition leads to response in some patients with head and neck squamous cell carcinoma (HNSCC). Robust biomarkers are lacking to date.

We analyzed viral status, gene expression signatures, mutational load and mutational signatures in whole exome and RNA-sequencing data of the HNSCC TCGA dataset (N = 496) and a validation set (DKTK MASTER cohort, N = 10). Public single-cell gene expression data from 17 HPV-negative HNSCC were separately re-analyzed.

Among HPV-negative HNSCC, APOBEC3-associated TCW motif mutations but not total single nucleotide variant burden were significantly associated with inflammation. APOBEC3-enriched HPV-negative HNSCC showed higher T-cell inflammation and immune checkpoint expression. Mutations in immune-evasion pathways were enriched in these tumors. APOBEC3B and 3C expression was identified in tumor cells and correlated with tumor inflammation.

We identified an APOBEC-enriched subgroup of HPV-negative HNSCC with a distinct immunogenic phenotype, potentially mediating response to immunotherapy.

## Introduction

Cancer is a disease of the genome in that cancer cells have acquired somatic variants that prove advantageous for their growth. These mutations lead to changes in affected proteins and eventually cellular transformation. The altered proteins can be recognized by the immune system through presentation of peptides by the major histocompatibility complex (MHC), which allows for eradication of the tumor. Immune evasion is therefore considered one of the hallmarks of cancer (Hanahan and Weinberg, 2011). Immune checkpoint inhibitors (ICI) are an effective treatment option in a subgroup of patients in several cancer types including head and neck squamous cell carcinoma (HNSCC) (Ferris et al., 2016). The presence of an interferon gamma inflamed gene expression signature (IFNG signature or T-cell inflamed phenotype) (Seiwert et al., 2016; Ayers et al., 2017), expression of immune checkpoint PD-L1 (Ferris et al., 2016) and tumor mutational burden are associated with response (Van Allen et al., 2015; Cristescu et al., 2018). However, effective predictive biomarkers to guide ICI treatment in the clinic are lacking to date.

HNSCC is a common cancer type worldwide. It is mainly caused by tobacco and alcohol consumption, as well as infection with the human papilloma virus (HPV) (Rieke et al., 2016). These two groups (HPV-positive and -negative) are distinct entities with different outcome and different tumor biology (Keck et al., 2015). A better responsiveness of HPV-associated tumors to ICI has been suggested by early clinical data (Seiwert et al. 2016) but not confirmed in other studies (Mehra et al., 2018, Ferris et al., 2016). Immune activation due to immunological “foreignness” in virally induced cancers is a potential rejection mechanism (Blank et al., 2016). Additionally, an intracellular anti-viral response mediated by the APOBEC3-family of proteins leads to the accumulation of mutations and tumorigenesis (Henderson et al, 2014). In several cancer types, APOBEC-mediated tumorigenesis is increasingly recognized as an important mechanism independent of viral infections (Burns et al., 2013; Venkatesan et al., 2018). APOBEC activity can be inferred from an analysis of mutational signatures in the tumor genome. A so-called TCW motif has been identified as an APOBEC-specific mutational signature (Alexandrov et al., 2013). The role of APOBEC-induced mutations in HPV-negative HNSCC and its association with immune activation is unclear. We analyzed mutational signatures to uncover presumed mechanisms driving tumor inflammation in HNSCC.

## Results

### Identification of inflammation-associated mutational signatures

Mutation as well as expression data from head and neck squamous cell carcinoma samples were downloaded from the Cancer Genome Atlas (N = 496). The presence of a T-cell inflamed microenvironment was assessed in the samples by analysis of a six-gene IFNG signature (Chow et al., 2016). Total count of single nucleotide variants (SNV) did not correlate significantly with the IFNG signature (Fig. 1a).

**Fig. 1a:**
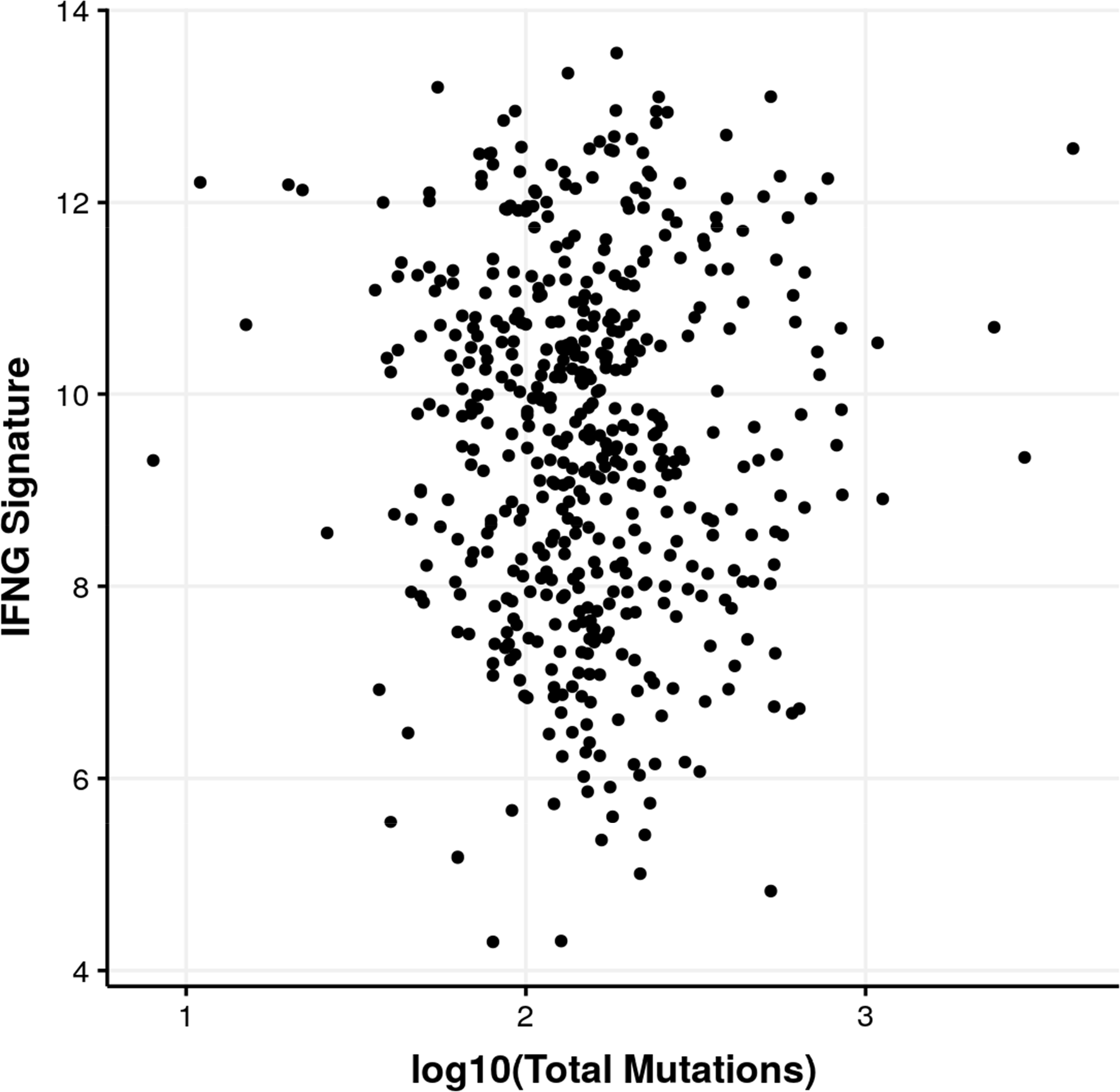
Relationship between total single nucleotide variant count and the six-gene IFNG signature. No significant correlation was found between these measures (R = −0.03 ± 0.08).

We then analyzed associations between the T-cell inflamed gene expression phenotype and mutational signatures. APOBEC3-associated mutations have been implicated before in HPV-positive HNSCC tumors (Henderson et al., 2014). These C>T and C>G mutations are sometimes referred to as TCW mutations, as they preferentially occur in TCA and TCT contexts (TCW motif). TCW mutations, where “W” corresponds to adenine or thymine, were significantly enriched in patients with high expression of the IFNG signature (Fig. 1b).

**Fig. 1b:**
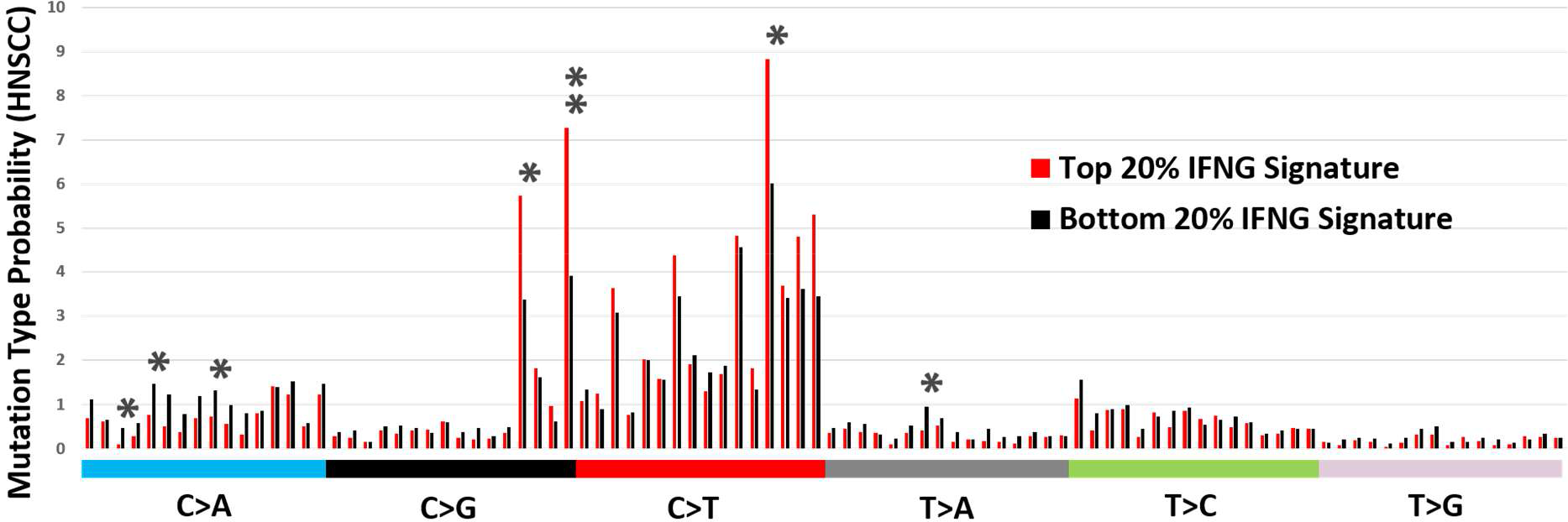
The frequency of base exchange motifs (e.g. C>A substitution with incorporation of the bases at the 5’ and 3’ end, thus allowing 96 potential mutation types) was compared between the patients with the highest and lowest IFNG signature within the TCGA cohort. The top 20% of inflamed cases showed a significant enrichment of variants in the APOBEC3-associated TCW context (* = p<0.05, ** = p<0.01, all p-values were Bonferroni corrected).

### Analysis of HPV-status on inflammation and mutational signatures

HNSCC consists of biologically distinct HPV-positive and -negative subgroups. We identified HPV-positive and -negative samples in the TCGA dataset (N = 64 and N = 432, respectively) by considering the amount of RNA-seq reads mapping to HPV types contained in the reference genome used by GDC (GRCh38.d1.vd1). Samples were considered HPV-positive if more than 3500 reads mapped to the respective top HPV contig (HPV types 16, 33, 35) with a median of 26916 reads. HPV-negative samples had a median of 1 read with a maximum of 85 reads. Results were checked for consistency against earlier results (Tang et al., 2013).

IFNG signature score, total SNV count, counts of TCW mutations and the ratio of the number of TCW mutations compared to the number of total mutations (“TCW ratio”) were assessed in both groups. Mutational load was significantly more pronounced in HPV-negative samples than in HPV-positive (Fig. 2a, p = 1.7 × 10^−4), whereas the IFNG signature was significantly higher in HPV-positive samples (Fig. 2b, p = 3.7 × 10^−5). Further, we compared the ratio of the number of TCW mutations / number of total mutations as a surrogate measure for APOBEC3 mutational activity, which was significantly higher in HPV-positive tumors (Fig 2c, p = 2.6 × 10^−3). We then analyzed the association between TCW mutations and inflammation in HPV-positive and -negative HNSCC and found a significant correlation only among HPV-negative HNSCC (Fig. 1c).

**Fig. 1c:**
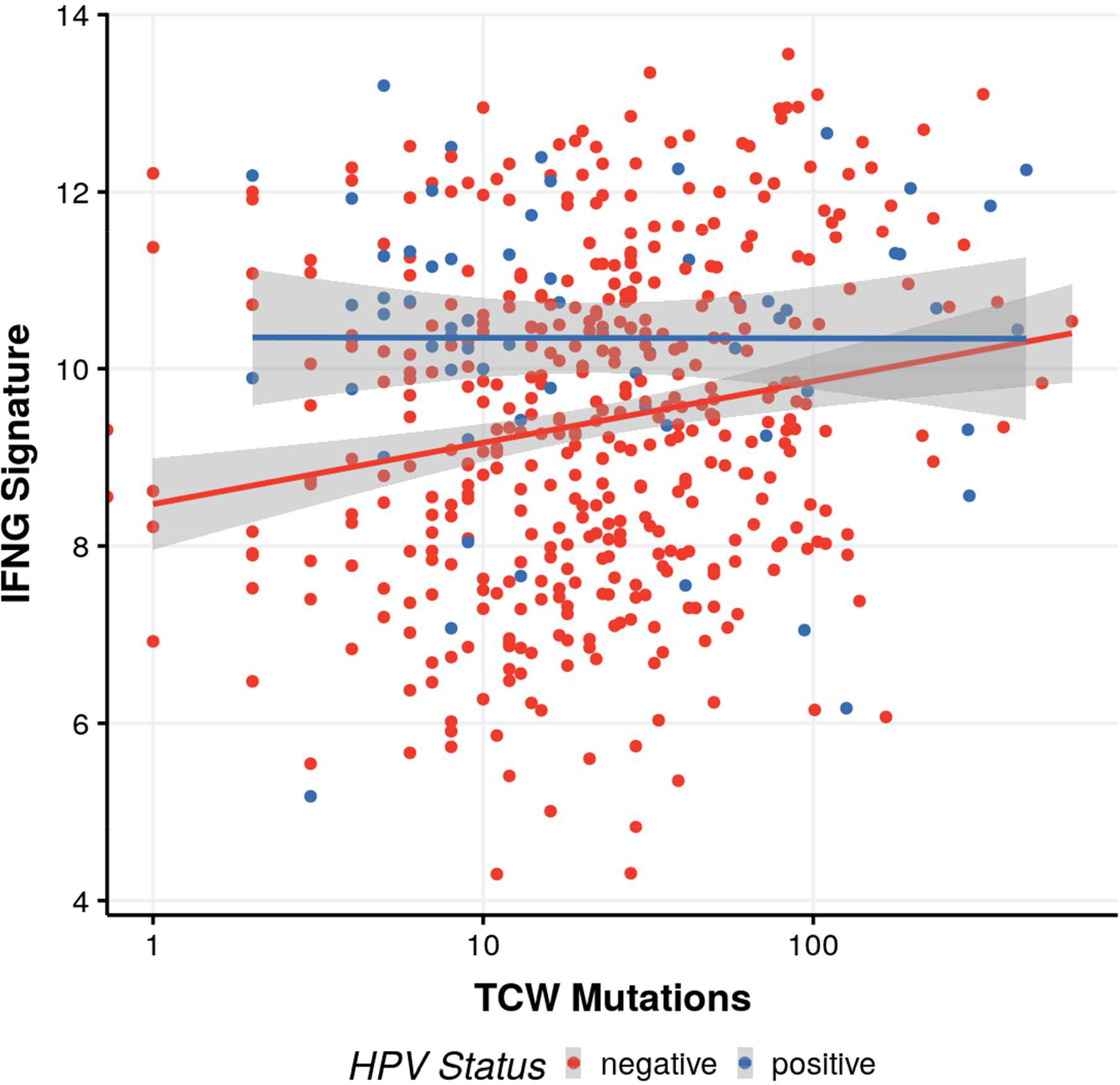
Relationship between the six-gene IFNG signature and the number of C>T and C>G mutations in TCW motifs. Colors indicate HPV-status (red: HPV-negative, n = 432, blue: HPV-positive, n = 64). A significant correlation between TCW mutations and the IFNG signature was identified in HPV negative cases (R = 0.18, p = 1 × 10^−4).

**Fig. 2a:**
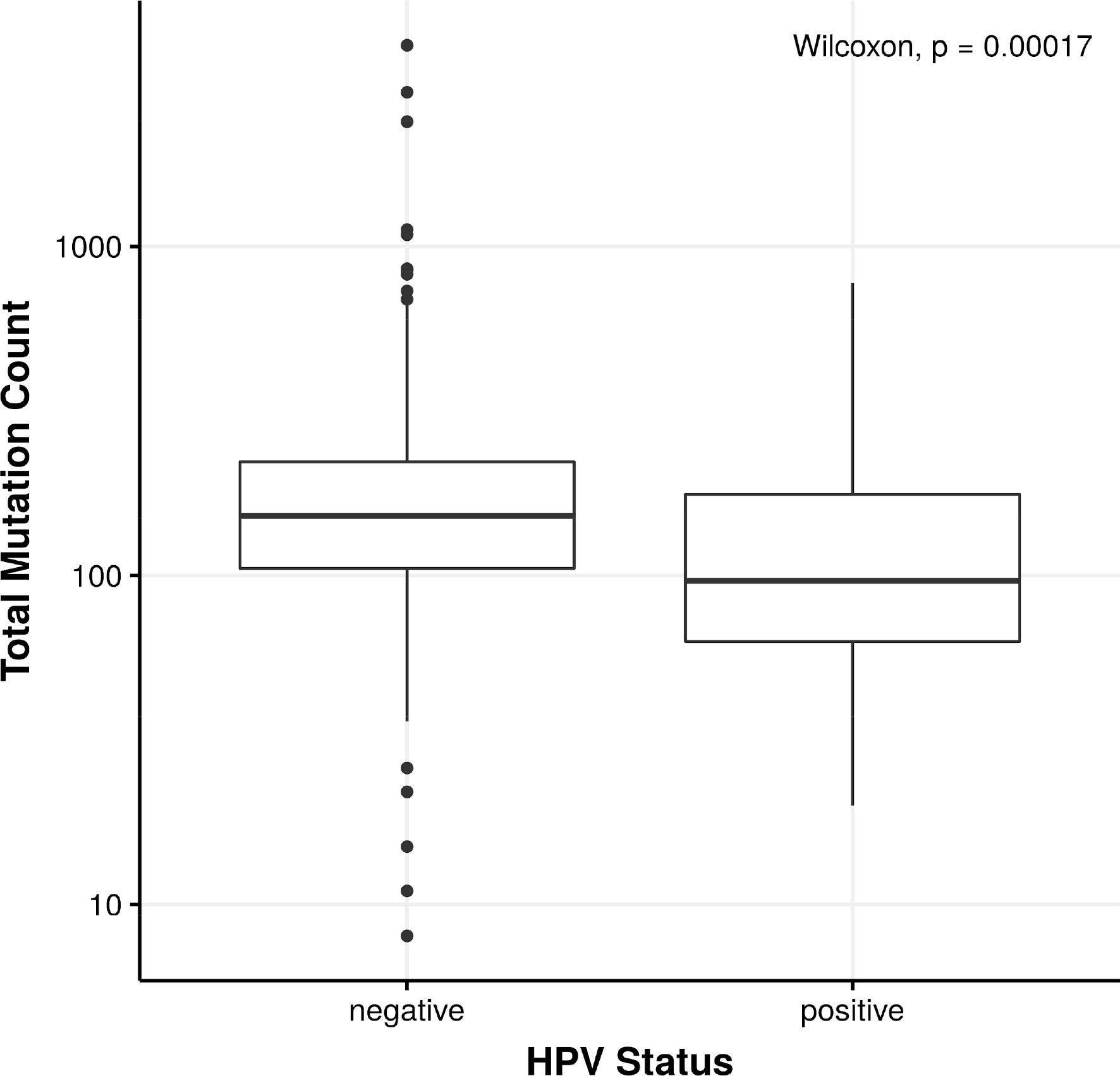
Boxplot of total single nucleotide variant count, grouped by HPV status. Total single nucleotide variant count was significantly higher in HPV -negative cases (p = 1.7 × 10^−4)

**Fig. 2b:**
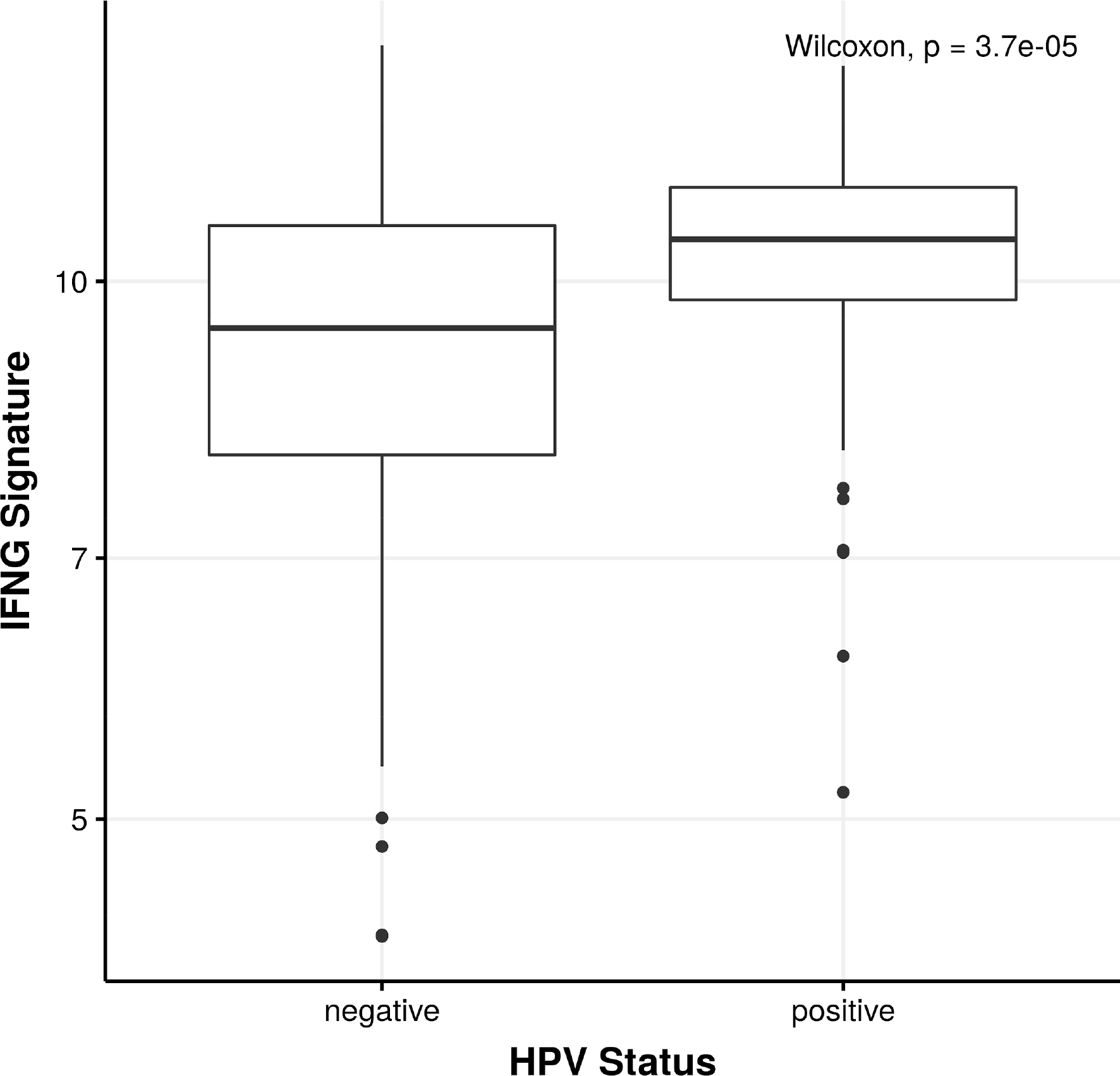
Boxplot of INFG signature score, grouped by HPV status. The six-gene IFNG signature score was significantly higher in HPV-positive samples (p = 3.7 × 10^−5).

**Fig. 2c:**
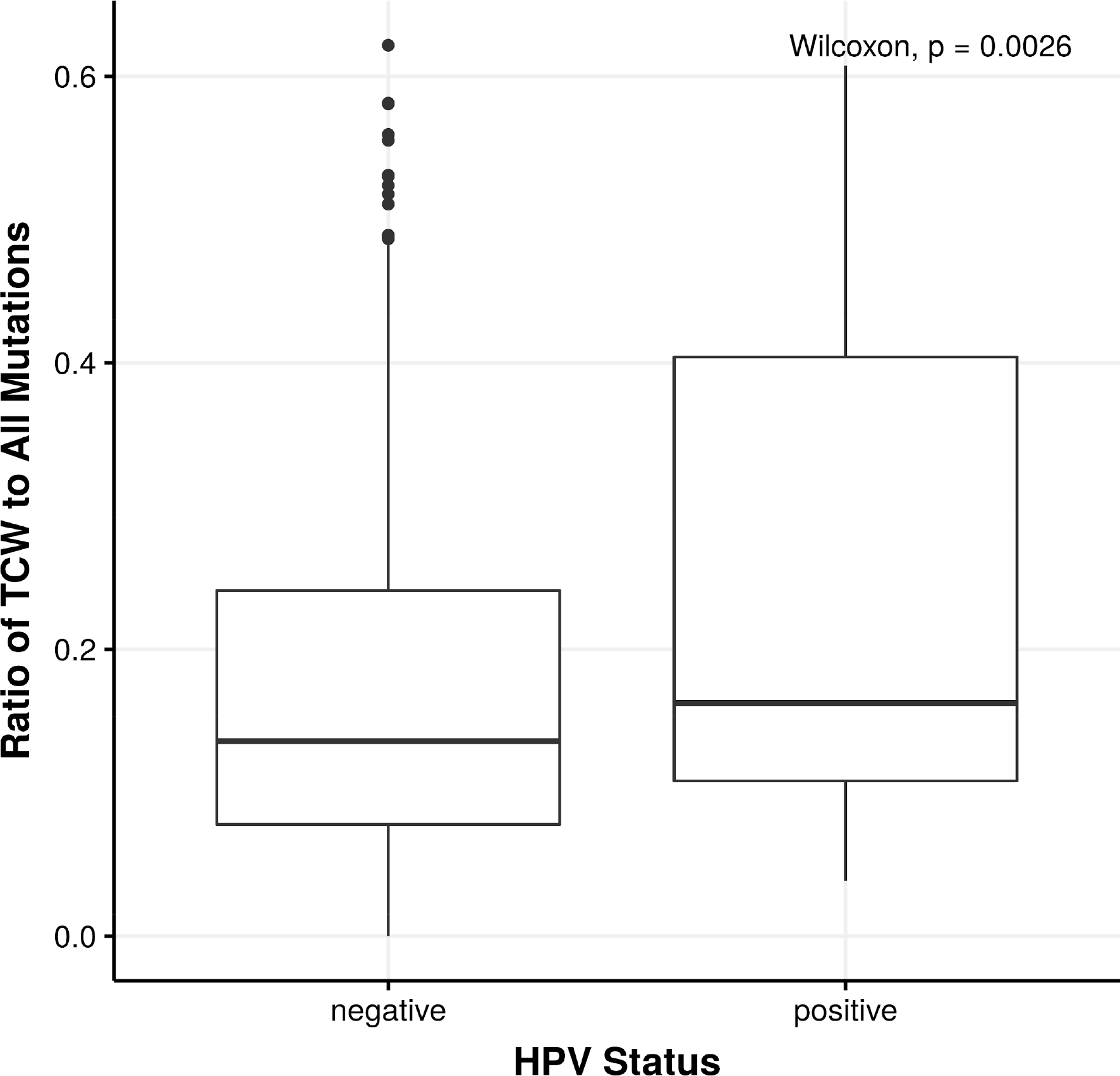
Boxplot of ratio of TCW variants to total single nucleotide variant count, grouped by HPV status. The TCW-ratio was significantly higher in HPV-positive samples (p = 2.6 × 10^− 3).

### Identification of an APOBEC-enriched HPV-negative subgroup

Since the association between APOBEC-induced mutations (TCW mutations) and the T-cell inflamed phenotype was restricted to HPV-negative samples, we grouped HPV-negative samples into APOBEC-enriched (N = 84) and APOBEC-negative (N = 348) cases. For each case, a Fisher test comparing the number of mutations in the TCW motif and the total number of cytosine mutations compared to their occurrences on chr1 of the human genome (Roberts et al. 2013) was used. P-values were Holm-Bonferroni corrected subsequently and all cases with p’ < 0.05 were labeled APOBEC-enriched.

We observed that the HPV-negative subgroup with an enrichment of putative APOBEC-induced mutations (HPV-negative, APOBEC-enriched) showed a significantly higher IFNG signature score compared to all other HPV-negative cases (HPV-negative, APOBEC-negative) (Fig. 3a). To exclude the possibility that this signal came from samples falsely classified as HPV-negative, we repeated the analysis by removing all HPV-negative cases with more than 5 reads mapping to any of the HPV contigs. Results did not change in a meaningful way.

**Fig. 3a:**
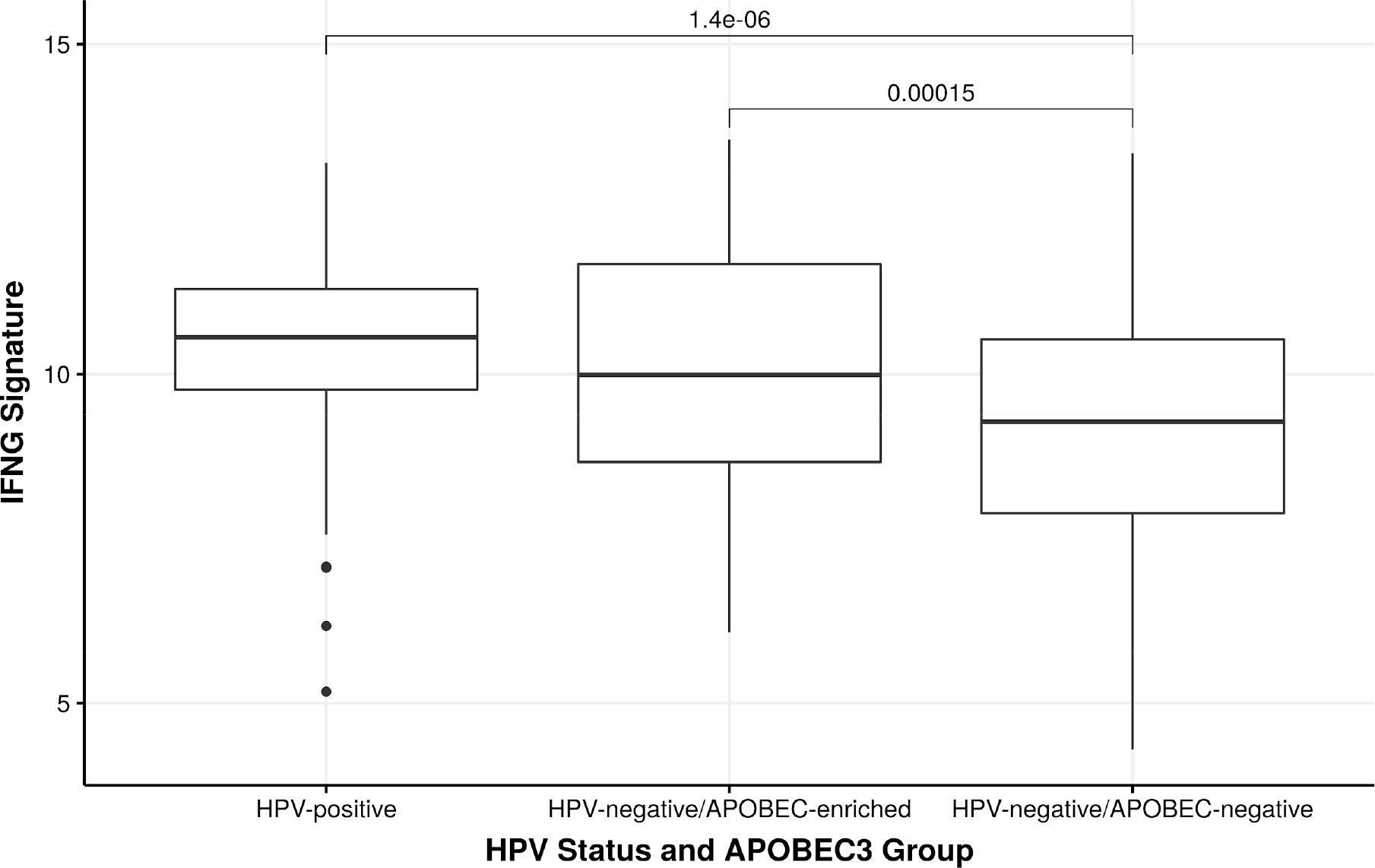
IFNG signature scores, grouped by HPV status and APOBEC enrichment for HPV-negative cases. HPV negative, APOBEC-enriched HNSCC showed a significantly higher IFNG signature than APOBEC-negative samples (p = 1.5 × 10^−4). No significant difference between HPV-positive and HPV-negative/APOBEC-enriched samples was found.

Further, differential expression of immune checkpoints was analyzed between groups. A significantly higher gene expression was identified for CD274 (PD-L1), CTLA4, LAG3, and PDCD1 (PD-1) in APOBEC-enriched cases. Only VTCN1 showed a significantly lower gene expression in APOBEC-enriched cases (Fig. 3b).

**Fig. 3b:**
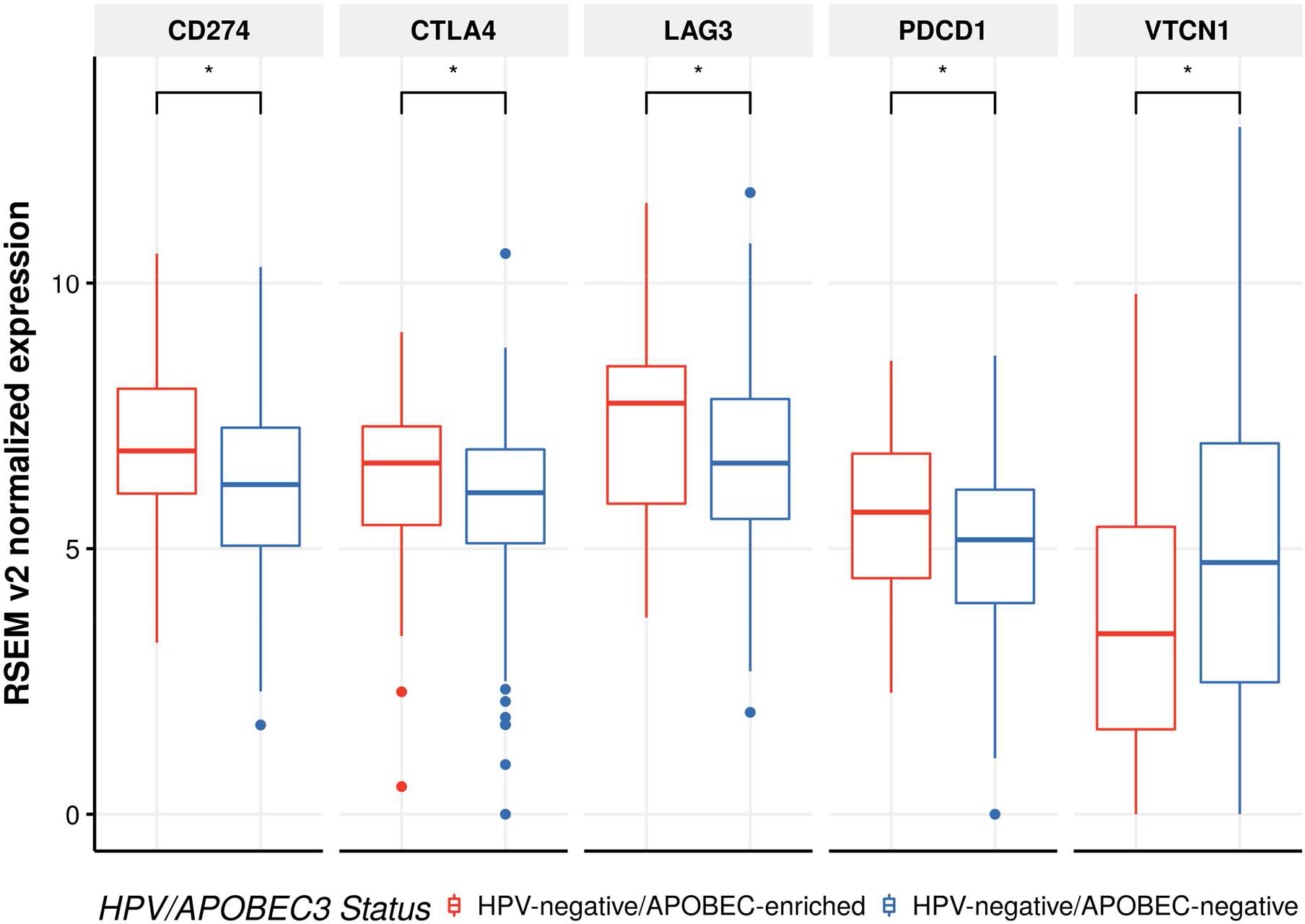
Gene expression of 5 immune checkpoints was significantly different between APOBEC-enriched and APOBEC-negative HPV-negative samples (p’ < 0.05 after Benjamini-Hochberg correction). All but VTCN1 showed significantly higher expression among APOBEC-enriched cases.

We further analyzed mutations in 554 genes which have been shown to be essential for cancer immunotherapy in a CRISPR assay (Patel et al., 2017, Supplementary Table 1). To identify enrichments of mutations in these genes, we used a Fisher exact test considering the number of cases in HPV-negative/APOBEC-enriched and HPV-negative/APOBEC-negative respectively, and the number of mutated genes from the aforementioned gene set in each group. The APOBEC-enriched subgroup showed significantly more variants with functional impact in immunotherapy-essential genes, as defined by Patel et al. (Table 1, p = 1.8 × 10^−4) (Patel et al., 2017). Among those genes, HLA-A showed the highest relative enrichment among APOBEC-enriched cases and remained significant after correcting for multiple testing (Suppl. Table 1).

**Table 1.** HPV-negative APOBEC-enriched and APOBEC-negative groups with counts of hits in gene set identified by Patel et al. Two sets of Fisher exact tests were carried out, first considering a gene mutated if any variant was found. Second, only mutations with putative functional impact (MODERATE, HIGH flags as returned by Jannovar (Jäger et al., 2014), e.g. missense or stop gain variants) were considered. Both times, the APOBEC-enriched group showed a significant enrichment for mutations in immunotherapy related genes compared to the APOBEC-negative group. Tests were re-done without variants in HLA genes to exclude possible false positive calls from variant calling.

|  | HPV-negative/<br>APOBEC-enriched | HPV-negative/<br>APOBEC-negative | P-Value |
| --- | --- | --- | --- |
| No. of Cases | 84 | 348 |  |
| No. of hits (collapsed to genes)/No. of cases | 7.6 (638 total) | 4.7 (1643 total) | 1.8E-4 |
| No. of hits with functional impact (collapsed to genes)/No. of cases | 5.7 (476 total) | 3.7 (1277 total) | 8.8E-4 |
| No. of hits (collapsed to genes), without HLA genes/No. of cases | 7.3 (617 total) | 4.7 (1623 total) | 4.1E-4 |
| No. of hits with functional impact (collapsed to genes), without HLA genes/No. of cases | 5.5 (461 total) | 3.6 (1259 total) | 1.6E-3 |

### APOBEC activation is an early event in some HNSCC

To identify temporal patterns of APOBEC activation during tumor evolution, we analyzed the variant allele frequency (VAF) of TCW mutations in HNSCC. Patients harboring significantly distinct variant allele fractions for TCW variants compared to all other variants of a given case were classified as early APOBEC3 activation, if the TCW variants had higher VAF, or as late activation, if they had lower overall VAF compared to other variants. Patients with no difference or too few variants were grouped as “no preference”. The false positive rate was controlled with the R package qvalue (Bass et al., 2018), using a threshold of 0.2.

Cases with early TCW variants exhibited a significantly higher IFNG signature score compared to late TCW activation (Fig. 4). We repeated the analysis for the TCGA cohorts of Urothelial Bladder Carcinoma (BLCA) and Lung Adenocarcinoma (LUAD).

**Fig. 4:**
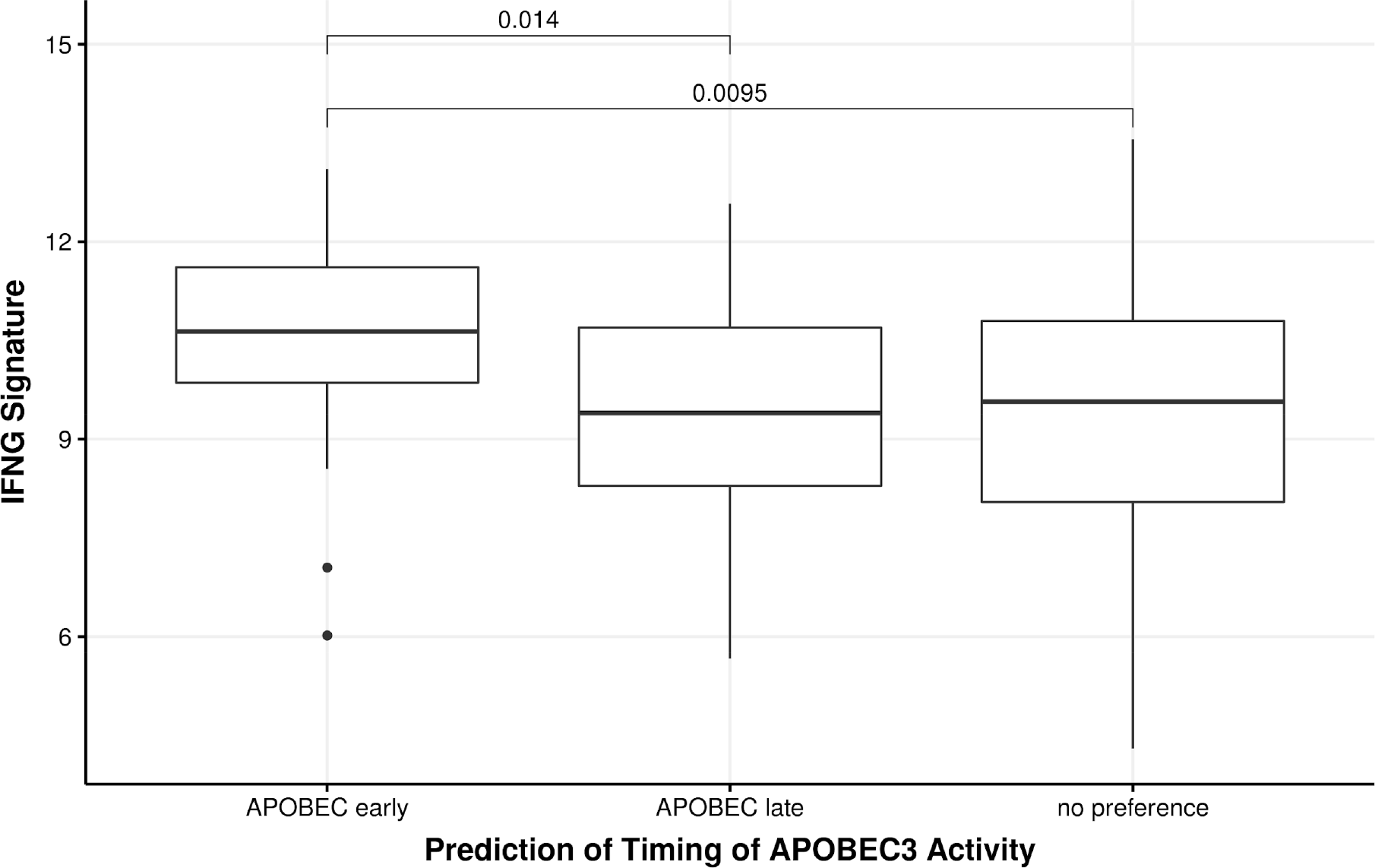
IFNG signature score in HPV-negative cases for each group of APOBEC3 activity timing. n(APOBEC early) = 15, n(APOBEC late) = 50, n(no preference) = 367. Inflammation was significantly higher in cases with putative early APOBEC3 activation (p < 0.05).

For BLCA, we did not observe any difference between the groups (Suppl. Fig 1). However, in LUAD, we observed the reverse effect. Cases with early APOBEC activation were found to exhibit lower inflammation scores than the other two groups (Suppl. Fig 2).

### Identification of APOBEC3B and APOBEC3C expression in HNSCC

Gene expression of APOBEC3 family genes was analyzed in the TCGA cohort. HPV-positive samples exhibited significantly higher APOBEC3 gene expression than HPV-negative samples. Among HPV-negative samples, the APOBEC-enriched subgroup showed significantly higher expression of APOBEC genes than the APOBEC-negative subgroup. Since bulk gene expression analyses do not differentiate between tumor and stroma, and immunohistochemistry of individual APOBEC proteins was not reproducible by our group (data not shown) and others (Venkatesan et al., 2018), we analyzed gene expression data in single cell transcriptome data of HPV-negative HNSCC (Puram et al., 2017, GSE103322).

Among those, APOBEC3B and APOBEC3C gene expression was detected in malignant cells (Fig. 5b-d), proving that the APOBEC3 expression signal that has been seen in bulk experiments does not exclusively originate from infiltrating immune cells, but also from the tumor itself.

**Fig. 5a:**
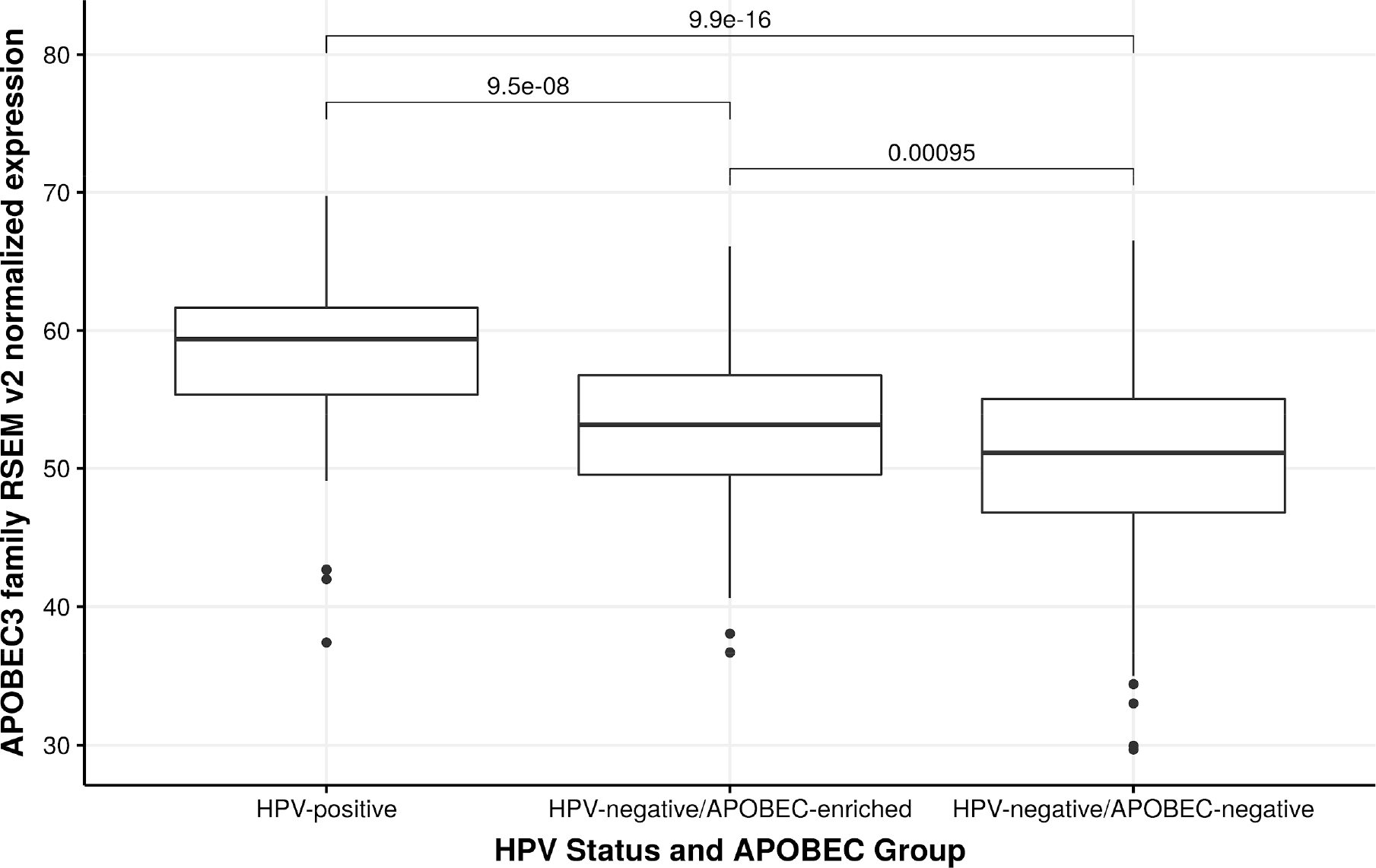
Aggregated expression of APOBEC3 family genes for the three groups HPV-positive, HPV-negative/APOBEC-enriched, HPV-negative/APOBEC-negative. HPV-positive cases show significantly higher APOBEC3 expression than HPV-negative cases. Among those, the APOBEC-enriched subgroup exhibits significantly higher APOBEC3 gene expression.

**Fig. 5b:**
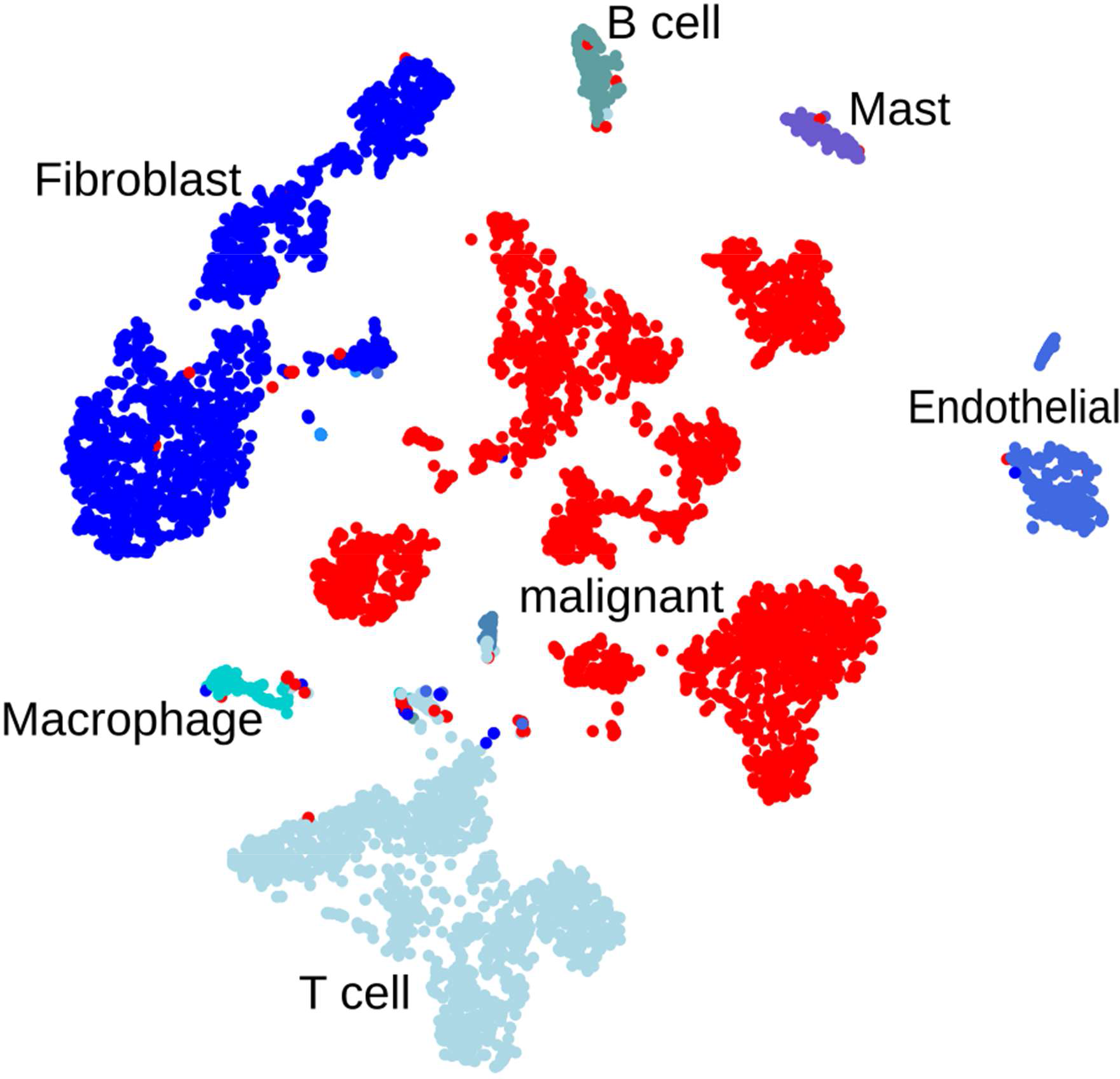
tSNE projection of all cells from 17 single-cell transcriptomics-profiled cases (Puram et al., 2017), grouped into cell types.

**Fig. 5c:**
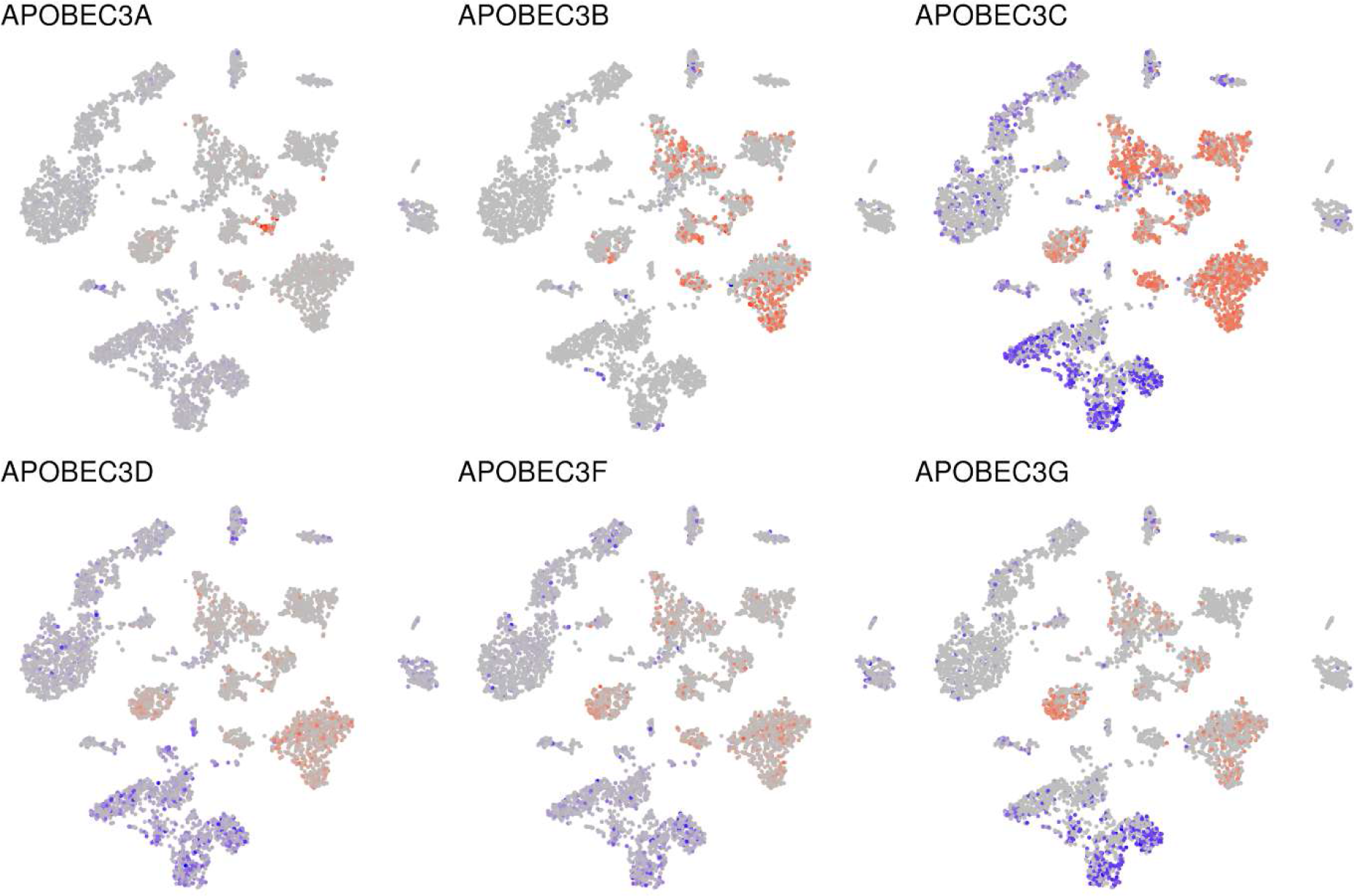
tSNE plots with projected expression of genes in the APOBEC3 family in 17 single cell data sets of HPV-negative cases (Puram et al., 2017). Biomarker-based groups of malignant (red) and non-malignant cells (blue). Expression strength indicated by color intensity, with grey indicating that no expression was detected.

**Fig. 5d:**
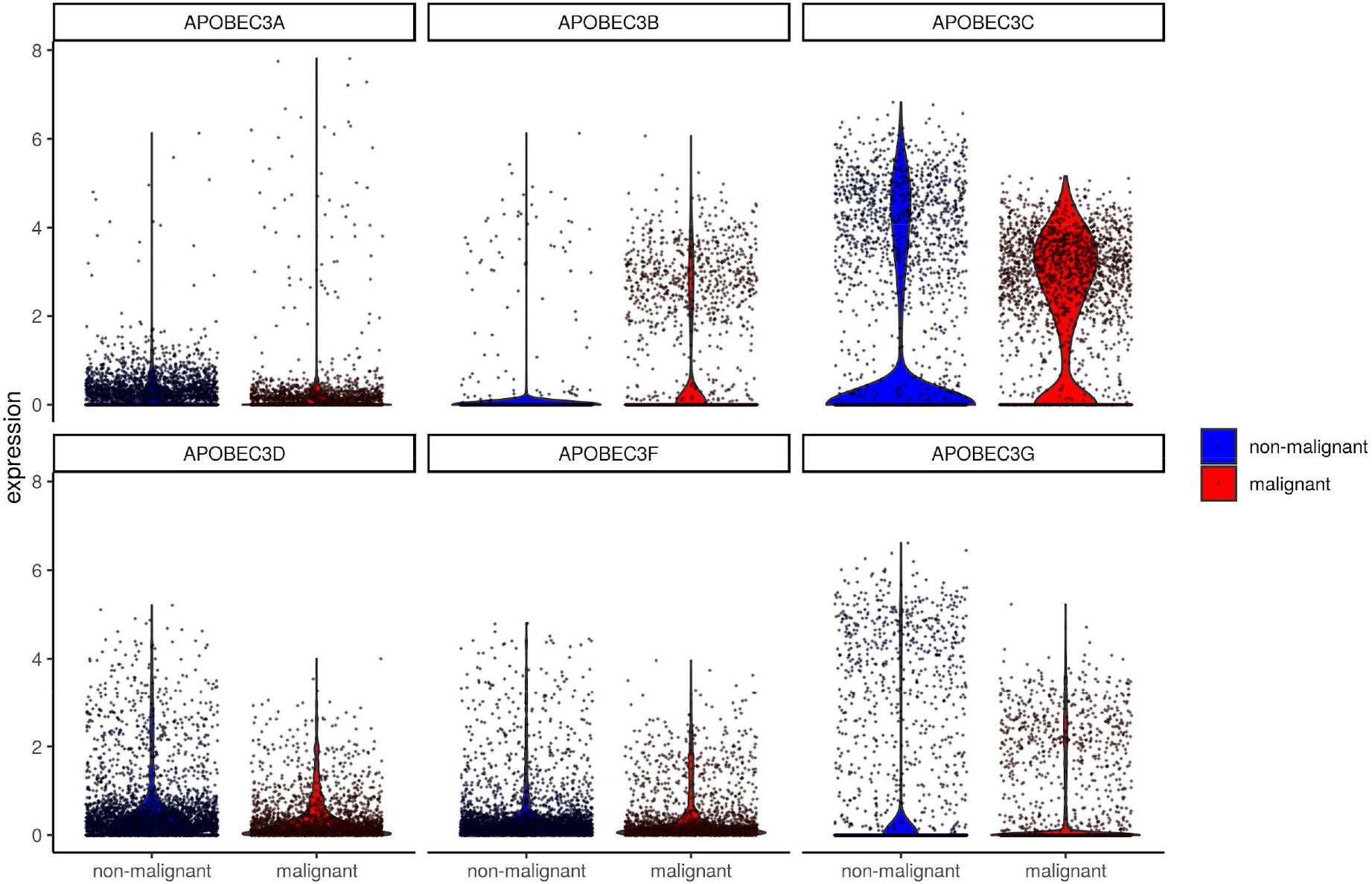
Violin plots of APOBEC3 gene expression between malignant (red) and non-malignant cells (blue). APOBEC3B and APOBEC3C gene expression was detected in tumor cells.

### Independent validation of findings

In an independent cohort of 10 HPV-negative cases of HNSCC, sequenced for the DKTK MASTER program from 6 cancer centers in Germany, we set out to validate these findings. We identified one HPV-negative patient with an APOBEC-enriched mutational signature. Again, we computed the IFNG signature score and compared it between the already defined groups (Fig. 6). The HPV-negative case with APOBEC3 activity (Fig. S3) showed the highest inflammation in this cohort.

**Fig. 6:**
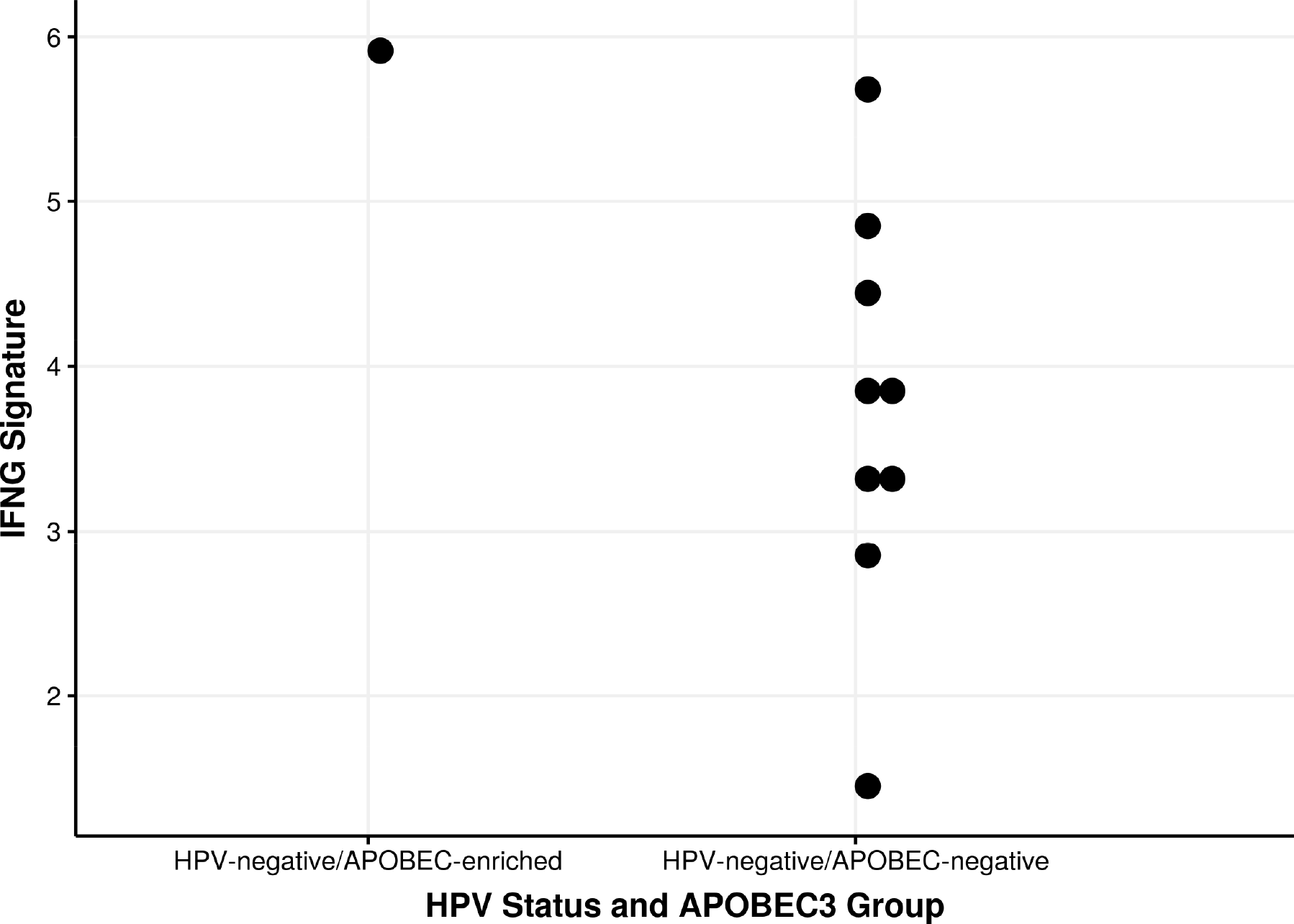
IFNG signature score in the head-and-neck DKTK Master cohort grouped by APOBEC status for all HPV-negative cases. The only identified HPV-negative/APOBEC-enriched sample harbored the highest IFNG gene expression signature score.

## Discussion

Immune checkpoint inhibition has become an important treatment option in HNSCC, providing a benefit in a subset of patients (Burtness et al., 2018). The predictive value of a T-cell inflamed phenotype, as defined by an IFNG expression signature, has been shown in HNSCC and other tumor types (Seiwert et al., 2016; Ayers et al., 2017). Additionally, a high mutational load is correlated with response to immune checkpoint inhibition (Van Allen et al., 2015). Yet, in our and other analyses, no clinically useful correlation between those two predictive markers was shown (Cristescu et al., 2018).

To better understand differential immune activation and evasion in HNSCC, we analyzed the relationship between different mutational signatures and inflammation in HNSCC. An APOBEC3 TCW mutational signature was significantly associated with a T-cell inflamed phenotype. This association was restricted to HPV-negative cases. We attribute this difference to the overall high impact of APOBEC-induced tumorigenesis in HPV-positive HNSCC (Henderson et al., 2014) and a generally higher level of IFNG activation in these tumors. The activation of APOBEC in non-virally-associated tumors has also been shown across cancer types (Alexandrov et al., 2013). This signature has been proposed to occur later in tumorigenesis and to induce branched evolution in lung cancer (Swanton et al., 2015). Additionally, an association has been shown between APOBEC and immune activation in lung cancer (Wang et al., 2018).

Our own analyses in HNSCC rather propose an early APOBEC activation in a subset of HPV-negative HNSCC with an immunogenic phenotype, thus proposing a differential oncogenic mechanism in this tumor type. It is currently unclear what drives this APOBEC activation. Previous analyses have suggested a link with single strand exposure and DNA repair defects (Taylor et al., 2013, Chen et al., 2014). It is also conceivable that short-term viral exposure induces APOBEC activation and carcinogenesis without genomic viral integration in some patients. A subgroup of HPV-negative oral squamous cell carcinoma patients in never-smokers, never-drinkers with high tumor inflammation has been described in the literature, accordingly and might correspond with our observation (Foy et al., 2017). Further research focusing on the impact of short-term viral exposure and APOBEC-activation or virus-independent mechanisms of APOBEC-activation, especially in this hard-to-treat subgroup, are of interest.

We were not able to immunohistochemically analyze APOBEC protein expression in HNSCC and HNSCC patient-derived xenograft models, a well-known problem with currently available APOBEC3 antibodies (Venkatesan et al., 2018). We therefore resorted to a re-analysis of publicly available single cell gene expression data. Doing so, we were able to circumvent the bias of measuring APOBEC3 activity in the tumor microenvironment (Leonard et al., 2016). Here, expression of APOBEC subtypes 3B and 3C was most prominent among malignant cells. APOBEC3B has also been identified in previous publications on APOBEC activation in cancer, including HNSCC (Burns et al., 2013; Wang et al., 2018), whereas the role of APOBEC3C remains less well defined.

It is currently unclear what causes the T-cell inflamed phenotype in APOBEC-induced cancers. We were able to show that these tumors, despite harboring the same overall mutational load, show a distinct immune escape, represented by an enrichment for mutations in immunotherapy-essential genes (such as HLA-A), as well as expression of immune checkpoint molecules. It is possible that APOBEC-induced mutations are more prone to detection by the immune system, due to their association with viral infections. Yet, our own analyses in APOBEC-induced cancers did not show an increase in bioinformatically predicted neo-antigens (data not shown). Therefore, the reasons for the different mechanisms of immune evasion in APOBEC-associated HPV-negative HNSCC are not known. Yet, these differences might also be relevant in urothelial (Glaser et al., 2018; Mullane et al., 2016) or lung cancer (Wang et al., 2018). Does this difference translate into differential response to immune checkpoint inhibition? Early studies suggest that APOBEC-associated tumors might indeed respond better to ICI therapy (Miao et al., 2018; Wang et al., 2018). These and our findings suggest to further investigate APOBEC-status and the use of mutational signatures from DNA sequencing as a predictive biomarker for immune checkpoint inhibition in HPV-negative HNSCC.

## Methods and Data

### Mutational load and APOBEC mutational signature

Somatic mutation data were downloaded from BROAD firehose for TCGA head-and-neck squamous cell cancer (2016/07/01). The MAF file was split into separate VCF files, one per TCGA sample. To annotate putative APOBEC induced mutations, we used the method described by Roberts et al. (2013), annotating C>T and C>G variants in TCW (TCA, TCT) motifs and their reverse complements, respectively. Further, we labeled all samples as “APOBEC enriched” if they showed a larger than expected amount of TCW vs C mutations when compared to a sample of the genome using fisher.test() from the R programming language and adjusting p-values with p.adjust().

### HPV status

HPV status was assigned based on the number of reads mapping to HPV genomes, which are now included as separate contigs in the newest bam files (genome release 38) available from GDC Portal. We set a cutoff of 3500 reads. Results were checked against the HPV expression signature described by (Buitrago-Pérez et al., 2009) and a derived reduced signature containing only gene CDKN2A and SYCP2 as well as prior results by Tang et al. (Tang et al., 2013) and TCGA clinical annotation for consistency.

### Expression data analysis

Expression data for sets of genes (“RNA Seq V2 RSEM”) was downloaded from the TCGA provisional cohort (Lawrence et al., 2015) from cbioPortal.org (Gao et al., 2013). IFNG signature was computed as the mean of the log2-transformed RSEM v2 expression values per sample.

### Single cell expression

This analysis was based on the digital expression matrix, holding the expression values of 23686 genes for 5902 cells of 17 tumor samples (GSE103322; Puram et al., 2017), together with a classification into malignant and various non-malignant cell types. Data were projected into tSNE coordinates using the standard Seurat workflow (Butler et al., 2018) and visualized using feature plots and violin plots.

### Independent validation of IFNG signature scores

Patients with advanced cancers, an ECOG performance status of 0-1 and an age < 50 years were eligible for enrollment in the DKTK-MASTER program across cancer centers in Germany. Whole-exome and RNA sequencing were performed on fresh-frozen tissues. From RNA-seq data for all cases, HPV status was predicted as described above. IFNG signature was computed as described above after generating transcript abundances with salmon (Patro et al., 2017) against ENSEMBL v75. The mapping of gene symbols used and their respective ENSEMBL ids is shown in Suppl. Table 2.

## Acknowledgements

This work was funded by a grant awarded by Berliner Krebsgesellschaft e.V. to DTR, FK and DB. DTR is a participant in the Berlin Institute of Health - Charité Clinical Scientist Program funded by the Charité - Universitätsmedizin Berlin and the Berlin Institute of Health. The results shown here are in part based upon data generated by the TCGA Research Network: http://cancergenome.nih.gov/. The DKTK MASTER trial is funded by the German Cancer Consortium (DKTK).

## Supplementary Material

**Suppl. Table 1:** Top hits to analysis in Table 1. Mutations in HLA-A showed the highest enrichment in APOBEC+ cases and remained significant after correcting for multiple testing (Benjamini Hochberg).

| Gene | p.adj | p | gene_mut_in_APOBEC + | gene_mut_in_APOBEC - |
| --- | --- | --- | --- | --- |
| HLA-A | 0,01153211 | 5.56E-06 | 20 | 16 |
| CASP8 | 0,23094184 | 2.23E-04 | 22 | 28 |
| HLA-B | 0,24745962 | 1.11E-03 | 11 | 10 |
| ODF2 | 0,24745962 | 1.31E-03 | 8 | 5 |
| SIK3 | 0,24745962 | 8.95E-04 | 7 | 3 |
| KDM4B | 0,24745962 | 4.20E-04 | 9 | 5 |

**Fig S1:**
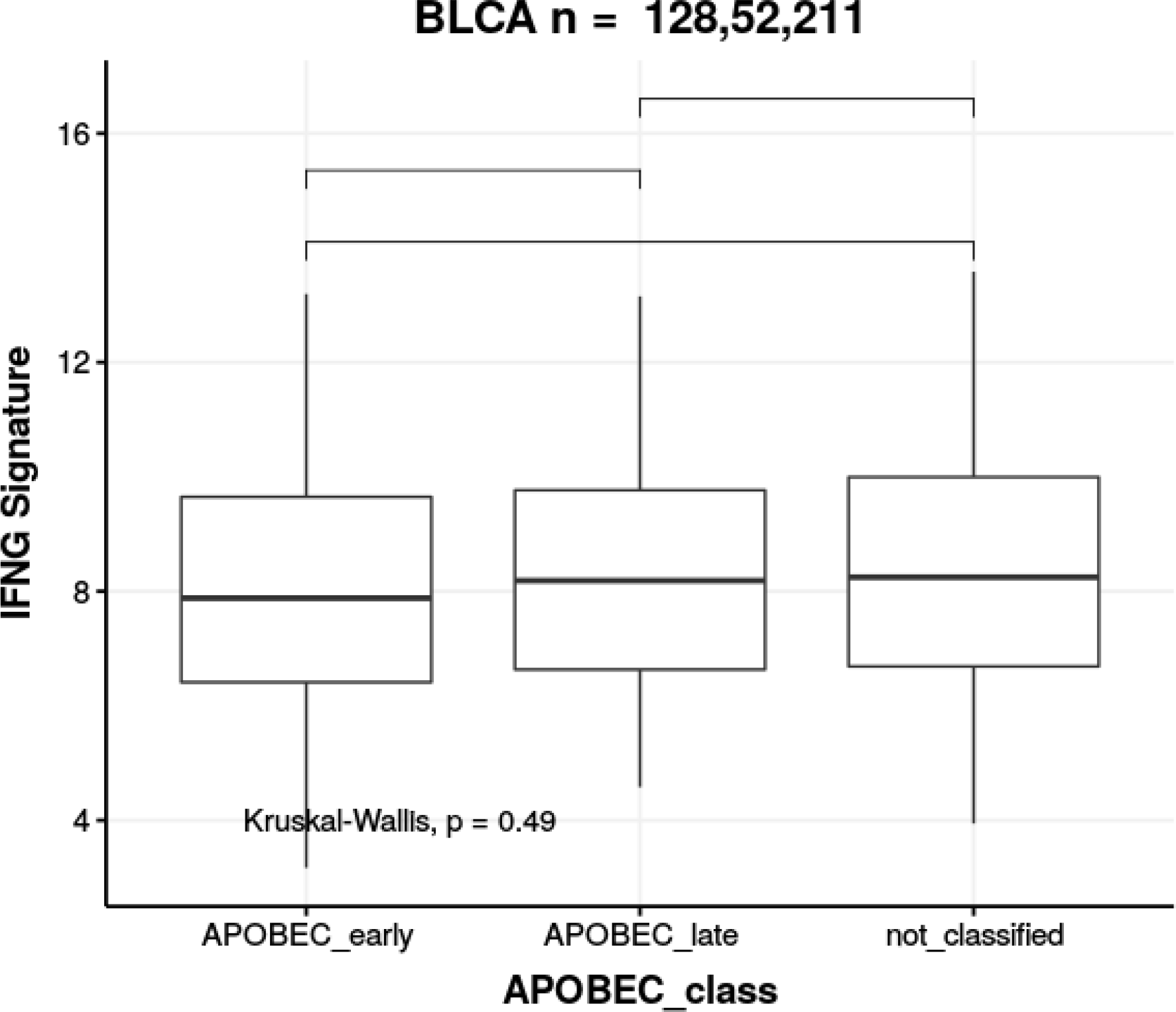
No significant differences were found between differential clonalities of TCW mutations among patients with bladder cancer.

**Fig. S2:**
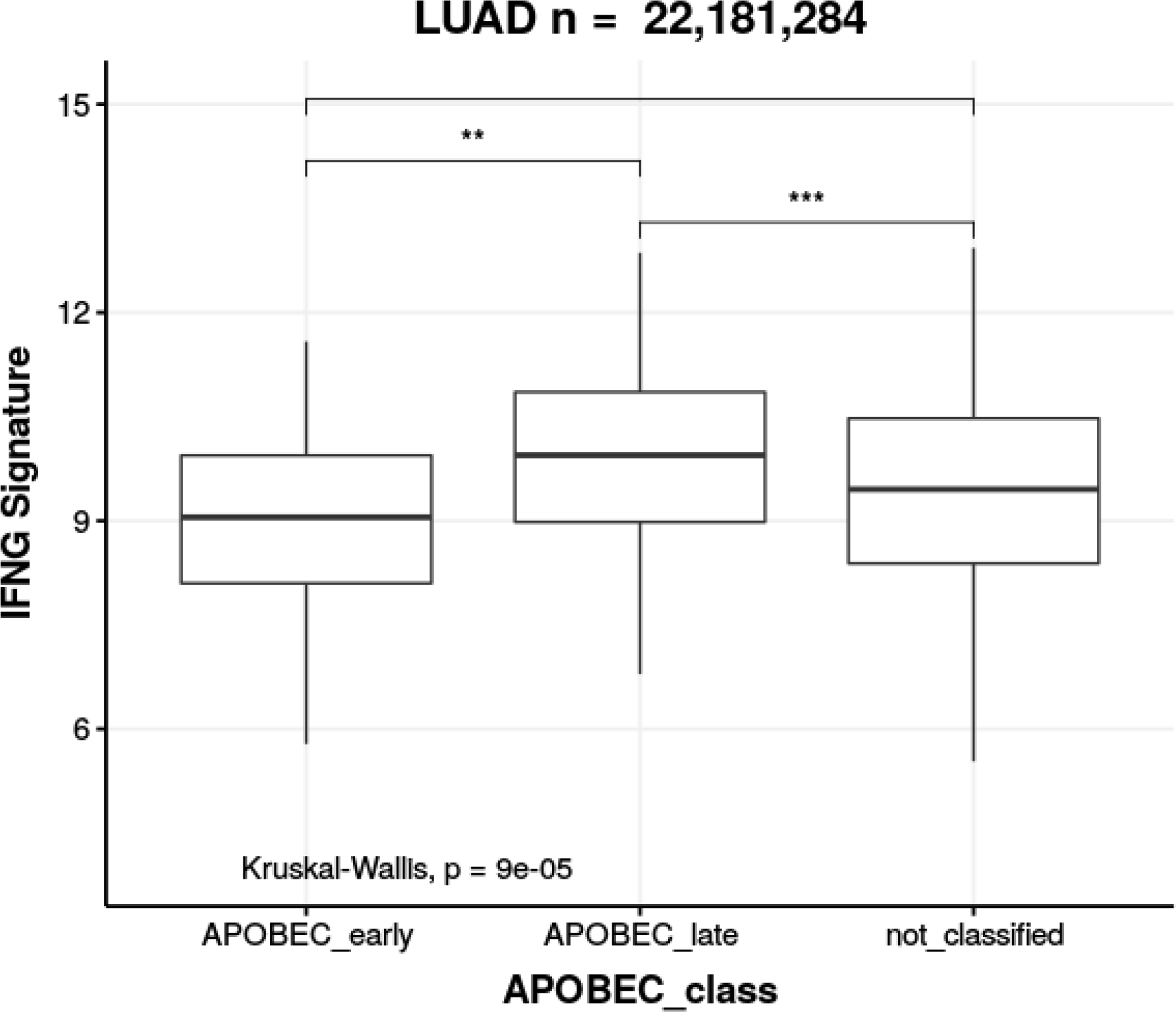
IFNG signature score in lung adenocarcinoma (LUAD) for each group of APOBEC3 activity timing. Inflammation scores are significantly lower in cases with putative early APOBEC3 activation.

**Fig S3:**
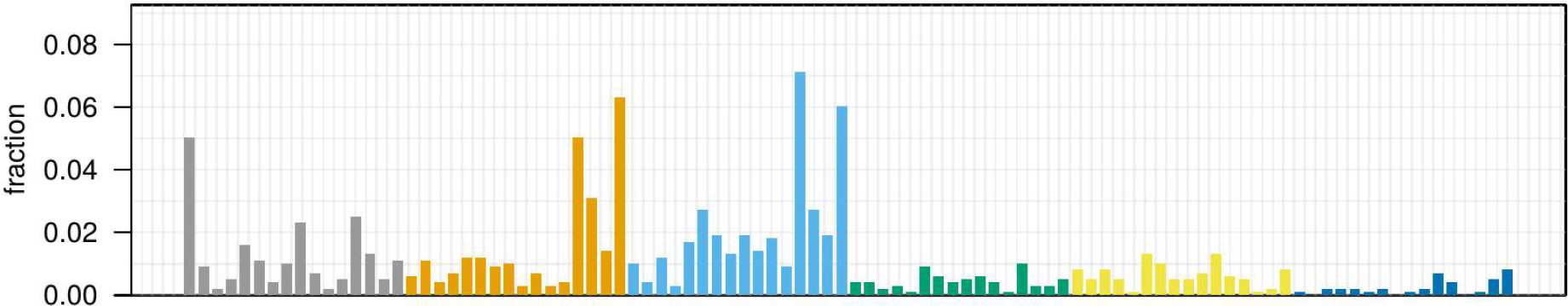
Mutational profile of the HPV-negative case with APOBEC activity.

**Suppl. Table 2:** Mapping of gene symbols to ENSEMBL ids for the IFNG signature (via ENSEMBL biomart)

| Gene stable ID | Gene name |
| --- | --- |
| ENSG00000111537 | IFNG |
| ENSG00000115415 | STAT1 |
| ENSG00000131203 | IDO1 |
| ENSG00000138755 | CXCL9 |
| ENSG00000169245 | CXCL10 |
| ENSG00000204287 | HLA-DRA |
| ENSG00000206308 | HLA-DRA |
| ENSG00000226260 | HLA-DRA |
| ENSG00000227993 | HLA-DRA |
| ENSG00000228987 | HLA-DRA |
| ENSG00000230726 | HLA-DRA |
| ENSG00000234794 | HLA-DRA |
| ENSG00000277263 | HLA-DRA |

